# LoxTnSeq: Random Transposon insertions combined with cre/lox recombination and counterselection to generate large random genome reductions

**DOI:** 10.1101/2020.05.25.114405

**Authors:** Daniel Shaw, Samuel Miravet-Verde, Carlos Pinero, Luis Serrano, Maria Lluch-Senar

**Affiliations:** Centre for Genomic Regulation (CRG), The Barcelona Institute of Science and Technology, Dr. Aiguader 88, Barcelona 08003, Spain; Pulmobiotics ltd, Dr. Aiguader 88, Barcelona 08003, Spain; Basic Sciences Department, Faculty of Medicine and Health Sciences Universitat Internacional de Catalunya, 08195 Sant Cugat del Vallès, Spain; Universitat Pompeu Fabra (UPF), Barcelona 08002, Spain; ICREA, Pg. Lluís Companys 23, Barcelona 08010, Spain

## Abstract

The removal of unwanted genetic material is a key aspect in many synthetic biology efforts, and often requires preliminary knowledge of which genomic regions are dispensable. Typically, these efforts are guided by transposon mutagenesis studies, coupled to deep-sequencing (TnSeq) to identify insertion points and gene essentiality. However, epistatic interactions can cause unforeseen changes in essentiality after the deletion of a gene, leading to the redundancy of these essentiality maps. Here, we present LoxTnSeq, a new methodology to generate and catalogue libraries of genome reduction mutants. LoxTnSeq combines random integration of lox sites by transposon mutagenesis, and the generation of mutants via cre recombinase, catalogued via deep-sequencing. When LoxTnSeq was applied to the naturally genome reduced bacterium *Mycoplasma pneumoniae*, we obtained a mutant pool containing 285 unique deletions. These deletions spanned from >50 bp to 28 Kb, which represent 21% of the total genome. LoxTnSeq also highlighted large regions of non-essential genes that could be removed simultaneously, and other non-essential regions that could not, providing a guide for future genome reductions.

## Introduction

One of the core principles behind synthetic biology is the rational engineering of an organism towards a specific application (D’Halluin and Ruiter, 2013; Esvelt and Wang, 2013; Mol et al., 2018). Genetic modifications that allow for the generation of new proteins are well documented and have myriad applications. Famous examples include modifying bacteria to allow us to study their functions in greater detail such as the production fluorescent proteins (Prasher et al., 1992), or genetically modified bacteria capable of producing molecules for human use such as insulin (Goeddel et al., 1979).

Live biotherapeutics have the potential to become key players in healthcare over the coming years. Engineered bacteria can be used as drug delivery systems, acting as a chassis from which therapeutic platforms can be plugged into to activate new functions (Ausländer et al., 2012; Chi et al., 2019; Claesen and Fischbach, 2015; Hörner et al., 2012; Vickers et al., 2010). A chassis will need to display growth characteristics and phenotypes that fulfil biosafety requirements, yet which may be foreign or counterproductive to cells within their original niche. For example, a limited ability to proliferate *in situ*, or to evade the immune system might be never selected for in the organism’s natural niche, but are features that might be of interest for a chassis. To illicit this change in phenotype, allowing the creation of an optimal chassis for a specific application, large-scale changes in genotype will have to occur. This will include both the removal of unwanted or unnecessary genomic regions, and the addition of new genes and functions (Folcher and Fussenegger, 2012; Ruder et al., 2011; Sung et al., 2016).

The loss of unwanted genetic regions has been shown to have desirable outcomes in bacterial engineering, with large scale genome reductions showing that the removal of superfluous genes can both increase production of a desired protein and develop beneficial characteristics for a cell (Chi et al., 2019; Sung et al., 2016; Vernyik et al., 2020). For example, sequential genome reduction has been used as an engineering tool in *Bacillus subtilis*, creating a mutant strain with a boosted capacity for production of heterologous proteins by systematically removing genes that hinder protein production, or divert energy and resources away from optimal protein production (Ara et al., 2007). Similarly, the removal of prophage elements in *Pseudomonas putida* resulted in a strain that demonstrated markedly higher tolerance to DNA damage (Martínez-García et al., 2015).

As more and more advanced engineering tools become available to researchers, the scope for genome engineering has similarly increased in scale (Annaluru et al., 2015; Cameron et al., 2014; Hsu et al., 2014). One of the grand long-term goals in systems and synthetic biology is the generation of a chassis with a minimal genome (Chi et al., 2019; Sung et al., 2016). While the definition of a true ‘minimal cell’ is almost impossible to define (Choe et al., 2016; Glass et al., 2017; Koonin, 2000), a general consensus has emerged around a cell with a reduced genome, capable of completing a specific task with as few superfluous functions as possible (Hutchison et al., 2016; Juhas et al., 2011; Zhang, 2010).

While naturally occurring ‘minimal’ bacteria do exist to some degree (Fadiel et al., 2007; Fraser et al., 1995; van Ham et al., 2003), with genomes as small as the one of *Mycoplasma genitalium* coding for only 470 genes (Himmelreich et al., 1997), the average gene complement for a bacterial cell is roughly 5000 proteins, though this can vary by two orders of magnitude between the extremes (Land et al., 2015). Comparative genomics studies, along with functional considerations give a hypothetical minimal genome consisting of 200-350 genes (Breuer et al., 2019; Dewall and Cheng, 2011; Gil et al., 2004; Glass et al., 2006; Koonin, 2000), so engineering a bacteria into a ‘minimal chassis’ requires large levels of genome reduction. On top of this, there is a further layer of small proteins and non-coding RNAs that are often overlooked, all of which have their own essentialities and interactions with the rest of the genome and cell (Lluch-Senar et al., 2015; Miravet-Verde et al., 2019).

Our understanding of epistatic networks, specifically clusters of genes that contribute directly or indirectly to a single phenotype (Phillips, 2008), and the complex web of interactions between gene circuits is far from complete (Otwinowski et al., 2018; Sailer and Harms, 2017; Weinreich et al., 2013). This has big implications for large-scale genome reduction projects, as knowledge gleaned from previous studies identifying non dispensable or essential (E) and dispensable or non-essential (NE) regions of DNA can become obsolete as soon as alterations are made to the genome. A good example of this can be found in the creation of the reduced-genome JCVI-Syn2.0 from *Mycoplasma mycoides* by the team at the J. Craig Venter Institute. After the creation of their landmark JCVI-Syn1.0 strain (Gibson et al., 2010), random transposon mutagenesis was performed on the organism to identify the NE genes. The genome was then divided into 8 sections, and each section had their NE genes removed independently, while the other 7 sections were kept intact. Despite removing only NE genes, only 1/8 configurations resulted in a viable cell (Hutchison et al., 2016).

A similar example of interdependence was seen when studying gene essentiality and metabolism in *M. pneumoniae* and *Mycoplasma agalactiae*. It was shown that in linear metabolic pathways producing an E metabolite, all genes were essential. However, when the E metabolite could be produced by two pathways, often both genes were classified as fitness (F) genes (Montero-Blay et al., 2020). Deletion of one pathway will make the genes in the other pathway become essential.

This is a key limitation in many genome reduction studies, as rationally designed genome reduction experiments often rely on data generated before genetic manipulations are introduced. As such, any *a priori* assumptions about gene essentiality can be potentially redundant after any reductions have occurred. Screening by transposon mutagenesis after every cycle of reductions can be applied (Hutchinson et al., 2016) but such screening efforts are both time and labour intensive, and relies on the basis that genes act alone, and not as a part of a larger network. However, the mutation or loss of certain genes has been shown to modulate the essentiality of others, known as “bypass of essentiality” (Ll, 2020). In the yeast *Schizosaccharomyces pombe*, 27% of E genes on chromosome II-L could be rendered NE in response to a different gene within the genome being deleted, mutated or overexpressed (Li et al., 2019).

This in turn demonstrates that reducing genomes based on *a priori* assumptions may not lead to the most optimised genome for the desired trait, as our knowledge on both how gene networks interact with each other on an epistatic level, and which genes are truly E or NE under any given configuration is still far from complete. On top of this, the essentiality of a gene is highly dependent on the environment the bacterium inhabits (Bloodworth et al., 2013; Sassetti et al., 2001), and a change in metabolic function may lead to a change in secreted or imported byproducts. This in turn could have knock-on effects on a cell’s microenvironment, causing certain genes to change their essentiality in unpredictable ways.

Previous studies have demonstrated that random deletions are feasible in bacteria as a genome streamlining methodology, notably in *Pseudomonas putida* (Leprince et al., 2012) and *Escherichia coli* (Vernyik et al., 2020). However both have been limited in either the ability to be scaled into a high-throughput screen or by the number of successful deletions reported.

Therefore, here we propose a new methodology, LoxTnSeq, to study the concept of genome reduction in an unbiased manner. Deletions are performed using the cre/lox system, consisting of two 34 bp lox sites which are acted upon by the cre recombinase. If both lox sites are in a *cis* orientation, the DNA between them is cirularised and removed from the genome (Ghosh and Van Duyne, 2002). We utilise mutant lox sites, specifically lox66 and lox71 (Albert et al., 1995), delivered via random transposon mutagenesis to remove large segments of genomic material via the action of the cre recombinase. As the cre can recognise and act upon these lox sites, we have designated them as ‘active’ here. To affect a deletion, the cre binds to the two active lox sites and recombines them, excising any DNA within them from the genome. As a result, a double mutant lox72 site is generated as a result of this recombination. This sequence is no longer recognised by the cre (Berzin et al., 2012; Van Duyne, 2015), and is therefore ‘inactive’. By combining a randomised genome deletion protocol with DNA-deep sequencing, we can identify the large putative reductions that are possible within a genome. This methodology allowed us to delete large sections of DNA without biasing the results by any potentially misguided *a priori* assumptions.

We have applied the methodology in the naturally genome reduced bacterium *M. pneumoniae*, considered a model for a minimal cell. However, this methodology could be applied to different bacterial systems in order to obtain different bacterial chassis, and thus to develop different applications in the synthetic biology field.

## Results

### Obtaining a high resolution library of lox mutants

Three different vectors (pMTnLox66Cm, pMTnLox71Tc and pMTnCreGm) were obtained, as described in Methods, to generate different libraries of transposon mutants. The plasmids here are derivatives of the Mini-pMTn4001 (Pour-El et al., 2002). Each plasmid contains a *Tn4001* transposase outside of the two inverted repeats flanking the cargo region, ensuring that the cargo is inserted without the transposon and is thus stable within the genome. The first library of transposon mutants was obtained by transforming the *M. pneumoniae* M129 strain with the pMTnLox66Cm vector. This vector inserted a lox66 site randomly in the genome. After selection with chloramphenicol, we obtained the first mutant pool.

The sequencing of the first round of transposition was analysed via HITS (High-throughput insertion tracking by deep sequencing) to properly assess the coverage of the first transformation. This methodology employs sequencing the DNA using an known oligo within the transposon that reads outwards, to identify where the transposon inserted within the genome (Gawronski et al., 2009). Insertions were mapped to *M. pneumoniae* (see Methods) revealing 355,319 unique insertions sites (Suppl. Table 1), which across the 816 Kb genome leads to a genome coverage of 43.5%, increasing to 64.1% when considering only known non-essential genes (Lluch-Senar et al., 2015). This represented an insertion every ~3 bases, an insertion frequency similar to what has been described in the latest transposition experiments using the same transposase and strain (Miravet-Verde et al., 2020). Figure 1 shows the insertion pattern of the pMTnLox66Cm transposon, along with the second pMTnLox71Tc transposon.

**Figure 1:**
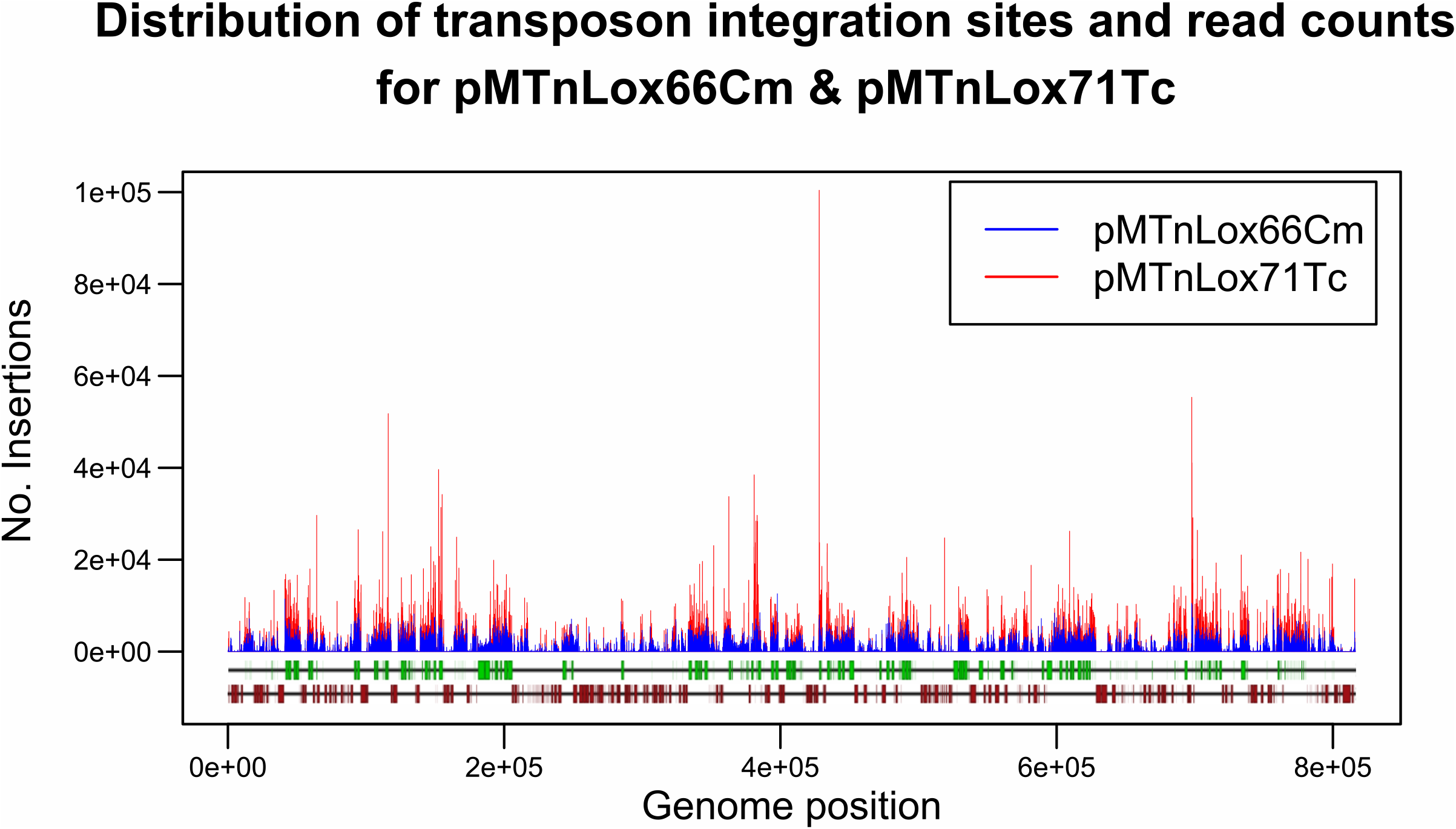
Transposon insertion points in the *M. pneumoniae* M129 genome, transformed in series with 1 pMole of the pMTnLox66Cm vector and subsequently 1 pMole of the pMTnLox71Tc, corresponding to the position of known essential (dark red boxes) and non-essential (green boxes) genes, as described by Lluch Senar et al. (<u>Lluch-Senar et al., 2015</u>).

### Creation of a pool of genome reduced mutants

This first pool of mutants was then transformed with the pMTnLox71Tc vector, and genomic DNA was isolated from the pool of surviving cells. After applying HITS and the same bioinformatic analysis procedure to this library, the number of unique insertions for the second transposon was 187,814. This leads to a genome coverage of 23% (1 insertion every ~4 bases), increasing to 38.2% for known non-essential genes. When considering together the samples covering the two transformation rounds, we accounted for a total of 387,963 unique insertions, corresponding to a 47.5% of genome coverage. Out of the 355,319 unique insertions found in pMTnLox66Cm, 155,170 (43.6%) were also recovered in pMTnLox71Tc leaving out 200,149 insertions positions (56,4%) found in the first sample but not detected after. On the other hand, sample pMTnLox71Tc presented 32,644 unique insertions (17.4%), with the remaining 155,170 shared insertions representing a higher portion of the total (82.6%) of the insertions found in the sample pMTnLox71Tc. In terms of read counts per insertion event, pMTnLox66Cm presented an average value of 250 reads per insertion increasing to 386 reads per insertion in pMTnLox71Tc. The locations and relative abundances for the two transformations are shown in Figure 1 and Suppl. Table 1.

We explored where the insertions were located at each transformation step by comparing the frequency of insertion for genome bins (Suppl. Table 2) and between the samples (Suppl. Table 3). Also, to detect any differential preference compared to a general transposon insertion protocol, we included in the comparative the essentiality reference samples for *M. pneumoniae* (Lluch-Senar et al., 2015). We observed a correlation between transformation steps of R^2^_genes_ =0.86 for genes and R^2^_bins_ =0.88 for bins. We also observed a better correlation between previous studies and pMTnLox66Cm (R^2^_genes_=0.88; R^2^_bins_=0.88) compared to pMTnLox71Tc (R^2^=0.81; R^2^_bins_=0.82).

We then explored the source of the differences found by calculating the fold-change of the first transformation over the reference samples. We observed that bins with fold-change > 2 (i.e. four times more insertions in pMTnLox66Cm compared to Lluch-Senar et al., 2015 dataset) were all in essential regions, thus with initial low frequency, and the observed enrichment was not significant because the difference in number of insertions was never higher than 50 insertions every 1kb bin. On the other hand, bins with fold-change < 2 (i.e. four times less insertions in pMTnLox66Cm) where all non-essential, and despite presenting less insertions when inserting lox sites, this bin regions where still non-essential in that condition (Suppl. Table 2). Considering these observations, we could assume the transformation with lox sites produces libraries comparable to a transposon sequencing regular experiment. When looking at the gene level, we observed a general trend of genes having less insertions in the second transformation compared to the first (Suppl. Table 4).

Both transposons utilise the same *Tn4001* transposase, and were grown under the same conditions (with the exception of selective antibiotic, see Methods), so we must assume that the loss of unique insertions sites can be a result of epistatic interactions from the first round of transposon insertion. This could be the presence of synthetic lethality pairs, or a case of selective pressure towards insertions that have the least fitness cost, due to the already compromised state of the genome. Despite the fewer unique inserts in the second round, there was still a coverage of 1 insertion every 4 bases on average, indicating a very large pool of potential deletions. In this way we generated a population carrying two lox sites, randomly distributed across the whole genome.

Importantly, transposon insertions take place randomly not only in terms of the chromosome position, but also in respect to their orientation. It is known that lox sites orientation is responsible from the action mode of cre (Van Duyne, 2015), catalyzing the excision of the DNA flanked by lox sites when they are in the same orientation, or the inversion of the region if lox sites are placed in opposite directions, as shown in Figure 2. Therefore, a population of cells carrying pMTnLox66Cm and pMTnLox71Tc insertions would undergo either genome reduction or inversion depending on the particular clone analyzed when subjected to the action of cre recombinase.

**Figure 2:**
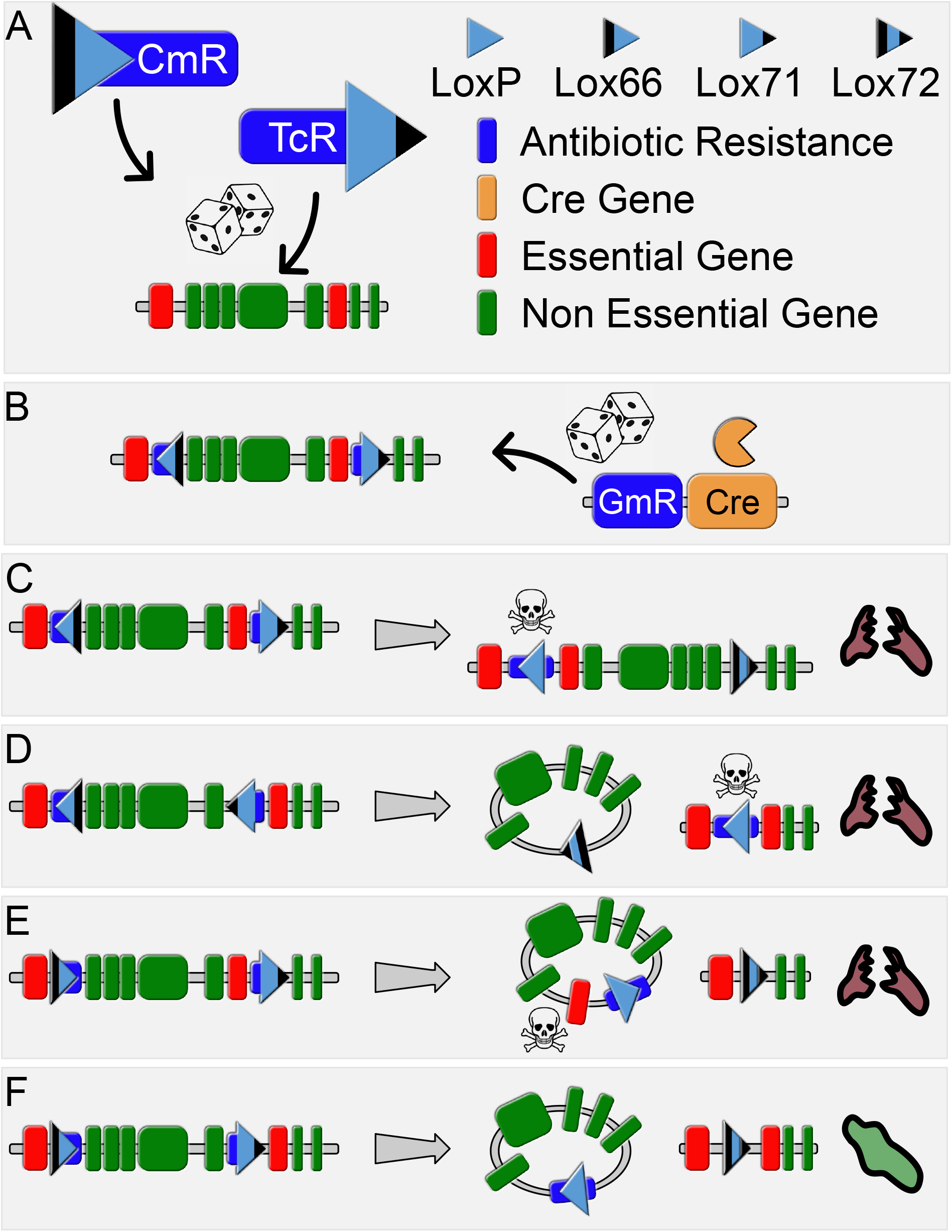
Schematic protocol, and interactions between mutant lox sites allowing for the selection of relevant deletion mutants. (A) Representation of the pMTnLox66Cm and pMTnLox71Tc transposons, along with dummy genomic DNA region and key. Both lox-containing transposons are integrated randomly into the genome (B) After transformation with the two lox site containing transposons, the third transposon pMTnCreGm containing the cre recombinase and gentamicin selective marker was added and randomly integrated into the genome. (C) Effect of the cre on lox sites integrating in *trans* orientation. The genomic DNA between the two lox sites is inverted, leaving a lox72 and loxP site. The active loxP in the presence of constitutively expressed cre is lethal in *M. pneumoniae*, and thus the cells containing an inversion are killed, indicated via a lysed purple cell. (D) Effect of the cre on lox sites integrating in *cis* orientation. Under the action of the cre, the genomic DNA between them is circularised and removed. Here, an active loxP remains within the genome, initiating a lethal phenotype, indicated via a lysed purple cell. (E) Effect of the cre on lox sites integrating in *cis* orientation, but encompassing an essential gene. The desired inactive lox72 site is left within the genome, but the loss of an essential gene causes cell death, indicated via a lysed purple cell. (F) Effect of the cre on lox sites integrating in *cis* orientation, containing no essential genes. An inactive lox72 is formed within the genome, impervious to the effects of the cre, and the non essential DNA is circularised and removed, with the green cell continuing to survive.

Cells containing both lox sites were then transformed with the pMTnCreGm transposon, without lox sites and containing the cre recombinase and a gene encoding gentamicin resistance. The cre expression was controlled by the constitutive p438 promoter (Montero-Blay et al., 2019; Pich et al., 2006), and was introduced via transposon to ensure high levels of expression. In previous work we demonstrated that the action of the cre on a single active lox site (i.e. one that can be recognised and acted upon by the cre, unlike lox72) in *M. pneumoniae* produces a lethal effect on par with a double stranded break in the DNA (Shaw, 2019). This was corroborated here, as surviving colonies were isolated and grown in media containing either gentamicin, or chloramphenicol and tetracycline, to assay which cells contained deletions and which cells contained inversions. 100/100 colonies picked grew in media containing gentamicin, indicating the cre/gentamicin transposon was present, yet 0/100 colonies grew in media with chloramphenicol and tetracycline, indicating 100% of the colonies contained a reduced genome and could no longer express the resistance genes on the lox transposons and that the inversions were fully removed, as shown in Figure 2.

### Identification of random deletions

Genomic DNA from the gentamycin-resistant pool of cells was sequenced via the circularisation protocol described in Methods. This methodology was chosen so that both sides of the deletion could be identified in a single read. Due to the random nature of the deletions, the only ‘known’ region of DNA to sequence from is the lox72 site, which is flanked by two ‘unknown’ genomic regions. Sequencing linear DNA would only reveal one side of the deletion per read, and it would then be impossible to pair the two reads from the same deletion, as all reads would start from the same point. By circularizing the DNA first, a single read can encompass both sides of the deletion. A total of 1,291,712 reads were recovered. Due to the random nature of the circularisation protocol, not all reads could reach both inverted repeats of the transposons and give accurate insertion points for each transposon. Therefore, to allow for as accurate mapping as possible for those reads that do not include both insertion points, the genome was split into 50 bp bins, and reads were grouped into these bins. The reads were then filtered by putative deletions that contained a known E gene (1365 unique reads, 83% of all unique reads), and those that did not (285 unique reads, 17% of all unique reads), according to Lluch-senar et al’s previous classification (Lluch-Senar et al., 2015). We found a background of deletions affecting E genes with few reads (41335 reads, 3% of total reads), which range in size from <50 bp to spanning over half the entire genome (See Figure 3), which probably are an artifact resulting from the circularization protocol for ultra-sequencing (see Methods). Those deletions affecting genomic regions with only NE genes were less in number, spanned smaller regions of the genome, with the longest continuously NE region being ≈30 Kb. However, they also could also contain circularisation artifacts. In total, they accounted for 1250337 reads (97% of total reads, see Figure 3). When excluding all deletions that contained an essential gene, 285 unique deletions were identified and mapped to the M129 genome (Suppl. Table 5).

**Figure 3:**
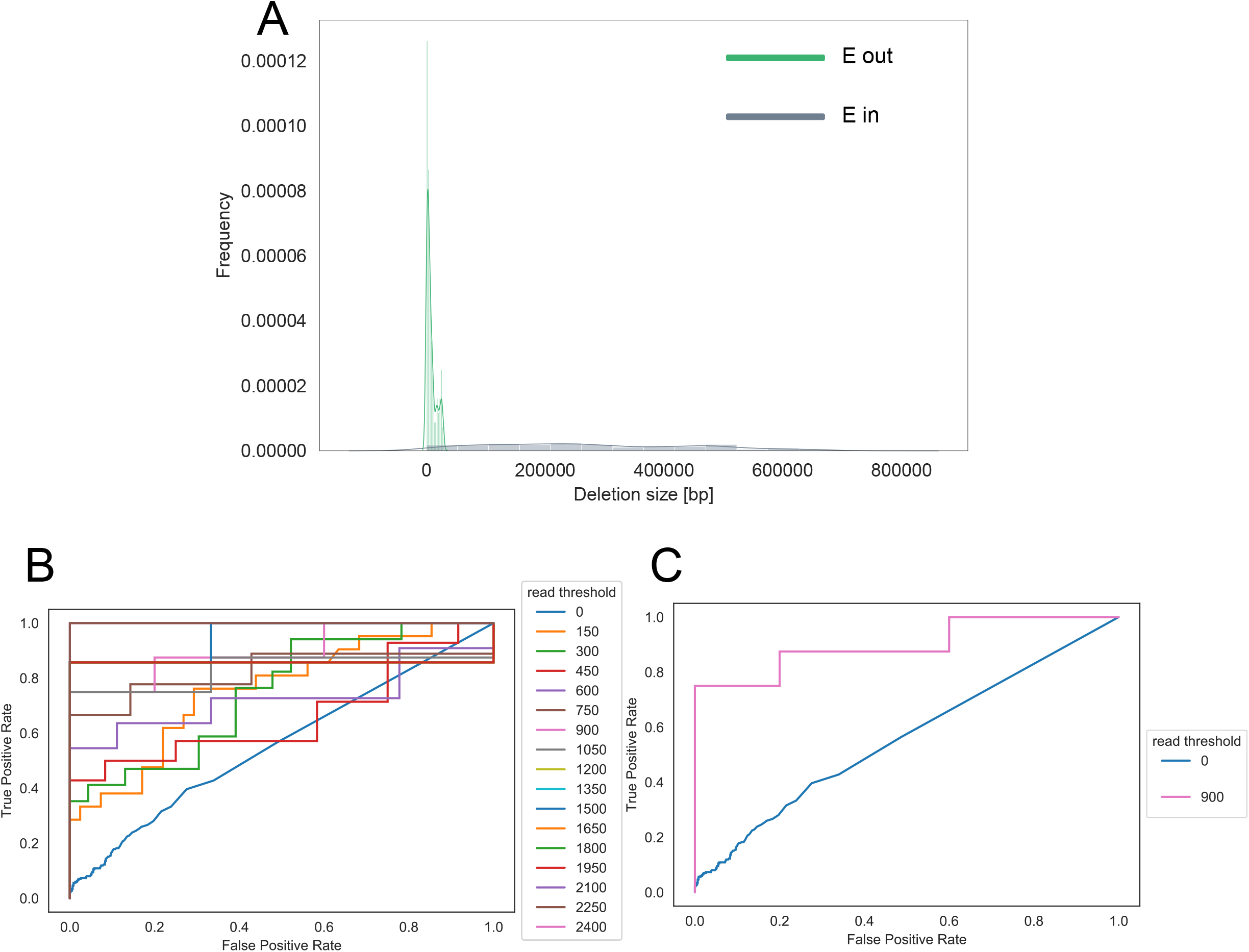
Distribution of sequencing data. (A) Relative abundances of the deletions that contain an essential gene (E in, blue) and those that do not (E out, green), based on the size of the deletion. (B) ROC curves assaying the false and true positive ratings of all deletions, depending on the number of reads. (C) ROC curve of our ‘Gold Standard’ delineation of 900 reads compared to the 0 read control.

Because the circularisation protocol involved fragmentation and re-ligation of the DNA, the possibility of spurious ligations occurs. Therefore, we needed a way of discerning which reported deletions were true and which were false positives, created via random DNA ligation. Using the read length as the threshold parameter, we performed a Receiver Operating Characteristic (ROC) curve approach to define high-confidence deletions (Figure 3B). This methodology allows the definition of a read count threshold maximizing the True Positive Rate (percentage of actual deletions properly detected) against the False Positive Rate (percentage of artifactual deletions wrongly detected as positive). Due to the fact that most of the positive and negative deletions were represented by very few reads, we observed that no discrimination could be performed without previous filtering based on read count. Thus, we iterated the process along with different pre-filtering and defined as best conditions to filter deletions with read count >900 that returned 8 deletions with a TPR >75% and no false positives (Figure 3C),. This set was classified as the ‘gold standard’ of deletions (Suppl. Table 5), as they had the highest level of reliability, and were mapped to the *M. pneumoniae* genome.

We then split the remaining 285 deletions that did not contain an essential gene via read count, those deletions that had greater than 10 reads, and those with less than 10 reads, which were also mapped to the *M. pneumoniae* genome, as shown in Figure 4A.

**Figure 4:**
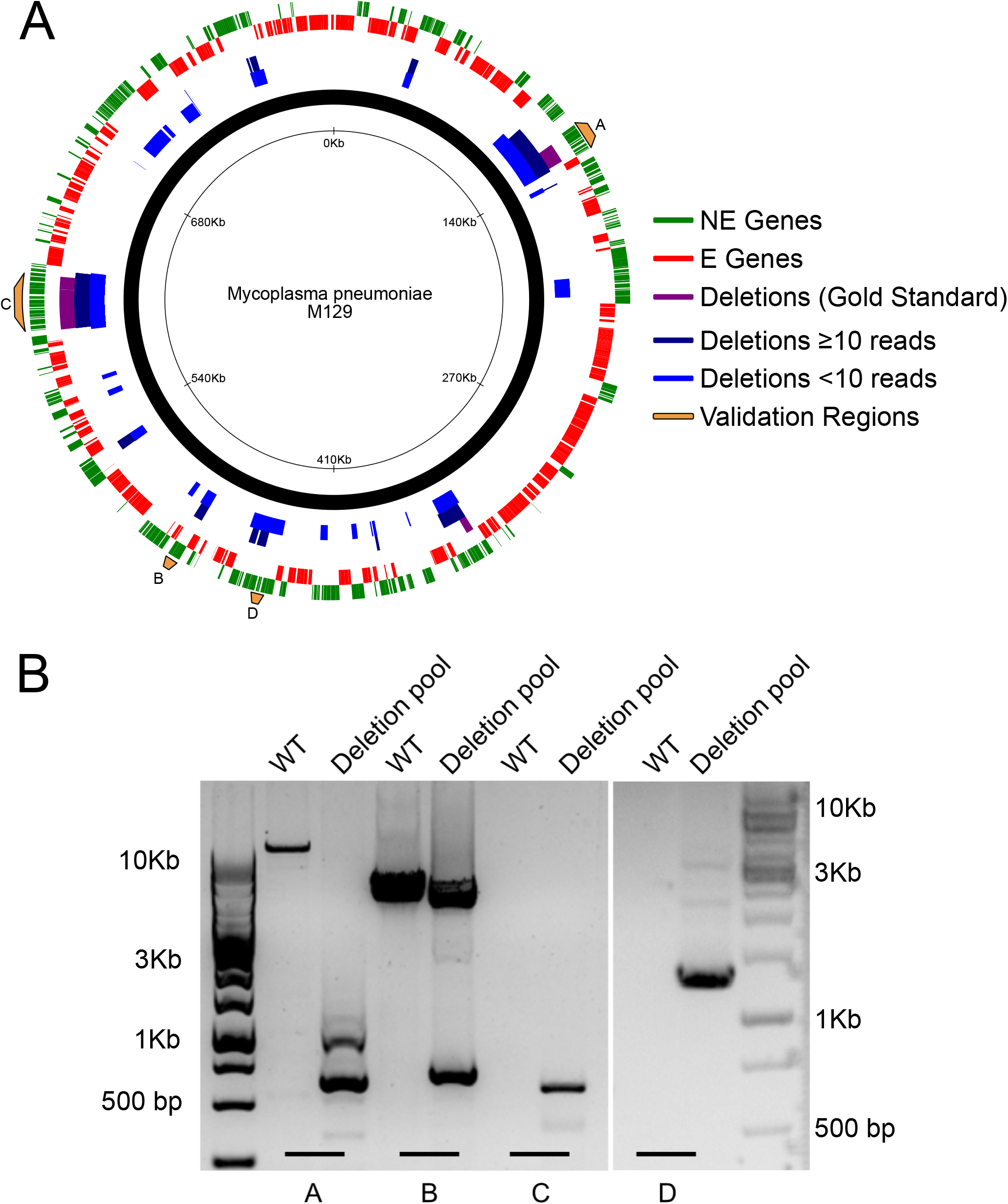
(A) Circos plot showing the locations of the non-essential genes (green, lane 1), essential genes (red, lane 2), deleted regions that were within the gold-standard (purple, lane 3), deleted regions with greater than or equal to 10 reads (dark blue, lane 4) and deleted regions with less than 10 reads (light blue, lane 5) relative to the *M. pneumoniae* M129 genome. Yellow regions indicate the areas amplified via PCR for validations. Created using CiVi (<u>Overmars et al., 2015</u>). (B) Validation of deletions via PCR. Gel electrophoresis of the four deletions candidates specified in the text above, with the WT on the left and deletions pool on the right in each condition. DNA primers were designed to amplify a 500-1500 bp region corresponding to the proposed deletion. Condition A amplified WT genomic DNA and genomic DNA isolated from the pool of deletion cells with oligos 11 & 12. Condition B amplified the same DNAs but using oligos 13 & 14, Condition C used the same DNA but with oligos 15 & 16, and condition D used the same DNAs but with oligos 17 & 18.

While the number of deletions within the gold standard is low, representing just 8 deletions (see Suppl. Table 4 and Figure 4A), they are all deletions spanning multiple genes, all of which are clearly well tolerated by the cell as they were highly represented via read numbers within the dataset. There are also many similarities between the sets, with similar deletions appearing in both highly and low read sets.

Despite the low number of ‘gold standard’ genes, we are confident that our sequencing method did not produce a large number of false-positives. As the data in Figure 3 shows, the vast majority of the reads generated (97%) indicated deleted regions that did not contain an essential gene, and the remaining 3% that did show an even distribution across the genome (see Suppl. Figure 1). Therefore, we feel confident applying the slightly less stringent test for fidelity, and simply allowing all the deletions that do not contain the removal of a known E gene.

Of the regions deleted, the largest was 28.7 Kb, the smallest <50 bp. 147 genes were deleted across the pool, accounting for 171.2 Kb (21% of the genome), with the vast majority of the functions unknown. Of the genes deleted, only 29 (19.7%) had an ascribed name and function. 139 genes were annotated as NE, with the remaining 8 classified as F genes, those genes whose loss is not lethal, but imparts a severe growth defect, according to the most recent essentiality data (Lluch-Senar et al., 2015). The full list of all deletions can be found in Suppl. Table 6, all deletions that did not contain an essential gene in Suppl. Table 5, and the genes deleted in Suppl. Table 7. Of the 259 genes that are classified as NE in *M. pneumoniae* (Lluch-Senar et al., 2015), we deleted 56%. The mean size of depletions comprising those NE genes was 7750 bp and median of 4750 bp, indicating the majority of genes deleted were removed as part of a larger region, not as single knockouts.

As validations, 4 regions were chosen from the list of putative deletions, labelled A, B, C, and D (see Figure 4A, yellow bars), with various characteristics. Region A contained seven NE genes (*mpn096* to *mpn102*), was ≈10 Kb in size, was the most highly represented deletion within the pool accounting for 964628 reads and the only validation within the gold standard. Region B represented a smaller region, containing four NE genes (*mpn397* to *mpn400*), a size of ≈5 Kb and covered by 593 reads. Region C represented one of the largest available deletions, and contained 19 NE genes and 1 F gene (*mpn493* to *mpn512*), with a deleted area of ≈25 Kb covered by 193 reads. Region D contained 6 NE genes (*mpn368* to *mpn373*) within 7.9 Kbs, but was represented by just one read in the dataset.

Figure 4B shows clearly that the putative deletion in all four regions were represented in the pool of deletions, but not found in the WT cells. There was no amplification in WT C due to its large size (25 Kb) There are also multiple deletions visible in the deletion amplifications in conditions A, B and D, which were expected from the sequencing data due to the multitude of different deletions in that area of the genome. The WT band is not visible in conditions C and D due to the PCR constraints in run length. The most prominent deletion band in the three conditions was isolated and sequenced, and each showed the requisite genomic regions intersected by the lox72 site within the transposon inverted repeats.

As the distribution of read counts was not uniform among the dataset, nor was the distribution of deletions uniform across the genome. There were clear hotspots present where multiple different deletions were found across the population, as shown by Figure 5.

**Figure 5:**
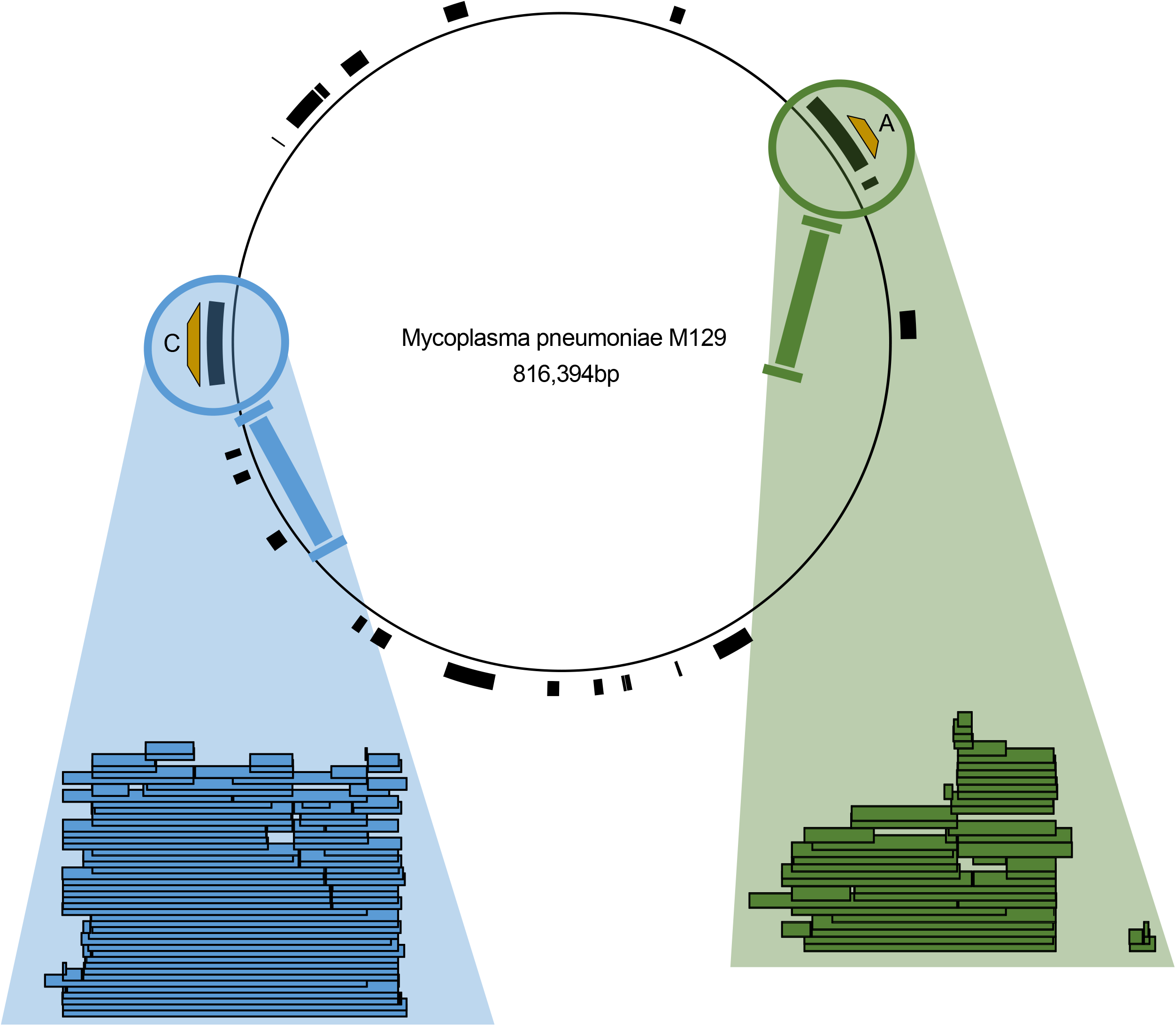
Location of all putative deletions with the M129 genome, with deleted regions shown as black bars. Magnifying glasses show regions with multiple deletions clustering in single regions, with each coloured bar representing a unique deletion within the dataset. Regions A and C from Figure 4A are indicated with the relevant yellow bars.

**Figure 6:**
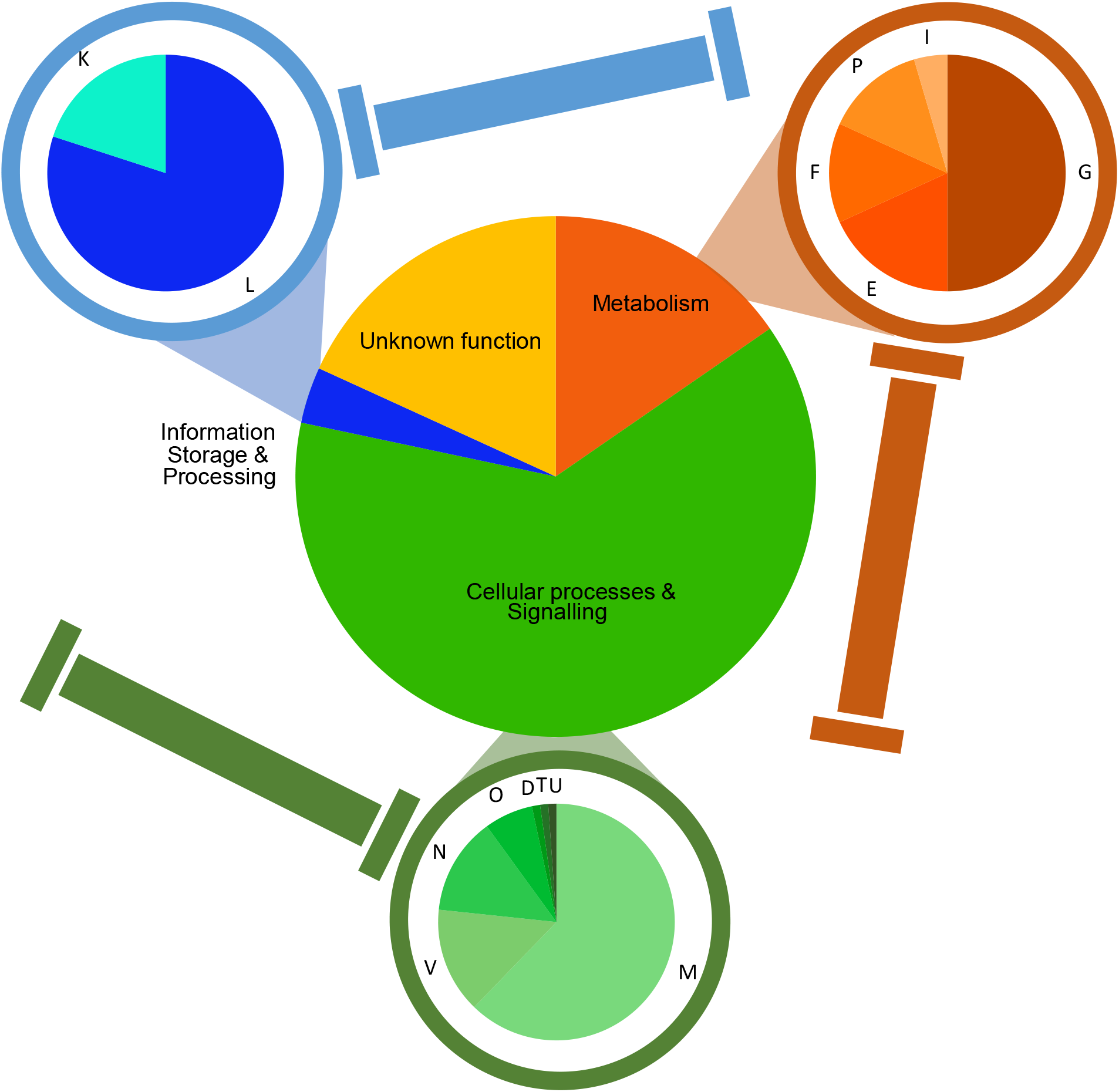
Breakdown of deleted genes via COG categories. [G] Carbohydrate transport and metabolism. [E] Amino acid transport and metabolism. [F] Nucleotide transport and metabolism. [P] Inorganic ion transport and metabolism. [I] Lipid transport and metabolism. [M] Cell wall/membrane/envelope biogenesis. [V] Defense mechanisms. [N] Cell motility. [O] Post-translational modification, protein turnover, chaperones. [D] Cell cycle control, cell division, chromosome partitioning. [T] Signal transduction mechanisms. [U] Intracellular trafficking, secretion, and vesicular transport. [L] Replication, recombination and repair. [K] Transcription.

The highlighted regions in Figure 5 show two of the strongest deletion hotspots, located at 120 Kb and 610 Kb regions, respectively. These regions show multiple overlapping unique deletions with slightly different lox insertions, across various sizes and configurations. This indicates that these regions are amenable to multiple different deletions, and there is little epistatic interaction between the genes present.

Contrary to this, Figure 4A shows a large NE region centered on the 200 Kb region of the chromosome. This region contains 13 NE genes (*mpn141* to *mpn153*), yet only 4 unique deletions were found within it, all of which were variations of a deletion from *mpn146* to *mpn152*. Despite the majority of the genes within this region having no known function, essential or otherwise, deletions within this region appear to be incompatible with cellular survival. The only known functions are linked to cytoadherence, specifically P1 adhesins (Nakane et al., 2011; Xiao et al., 2015). The canonical P1 adhesin (*mpn141*) was not deleted, nor were any of the other major adhesin genes *hmw1 (mpn447), hmw2 (mpn310) and hmw3* (*mpn452*). However, 15/22 adherence proteins were deleted across the population, so why this section is more essential than the others is unclear.

Looking at the functions of the deleted genes, they were grouped by COG category (Tatusov et al., 2000). The most commonly deleted functional category was M, genes involved in the composition and biogenesis of the cell membrane, accounting for 39% of deleted genes. Following this was the genes of unknown function (COG category S), with 18%. In total, deleted genes involved in metabolism (COG categories E, F, G, I and P) composed 15% of all deleted genes, information storage & processing genes (COG categories K and L) composed 3%, genes of unknown function 18% and genes involved cellular processes and signalling (COG categories D, M, N, O, T, U and V) 63% (see Suppl. Table 8 for full breakdown).

## Discussion

The cre/lox system has been used as a deletion system on many occasions, due to its ability to act in both prokaryotic and eukaryotic cells. In addition, the usage of mutant lox sites to facilitate deletions that result in an inactive lox has also been demonstrated in bacteria, such as being used to knock out single genes in series (Pan et al., 2011), or to knock out large but targeted genome region using either targetrons (Cerisy et al., 2019) or via recombineering (Xin et al., 2018).

A similar approach to this, using s transposon delivered FLP-FRT system instead of the cre/lox, was undertaken in *Pseudomonas putida*, creating large random deletions within the genome (Leprince et al., 2012). There, the authors succeeded in performing cyclical deletions within the genome, however there were three main caveats that our methodology aims to improve upon. First, their system is not self-selective for deletions. Individual mutants need to be screened manually for a loss of the two antibiotic resistances found in the transposons which would indicate a deletion has taken place. This precludes its use as high-throughput assay for surveying all possible deletions. Secondly, the efficacy of transformation was low, with only 255 independent insertions recorded for the second transposon insertion. This drastically lowers the utility in finding all possible deletions. Finally, while their system does allow for multiple rounds of deletion, this is limited by the usage of the FLP-FRT system. After a successful deletion has occurred, the system still contains a FRT site within the genome. Any subsequent insertions of a new FRT will utilise this pre-existing genomic FRT to create the deletion. Therefore, all subsequent deletions are limited to the areas directly adjacent to the original. While our system is not currently optimised for multiple round of deletion, the creation of the lox72 site via our methodology gives the potential for new deletions to attempted across the entire genome, as the lox72 is not recognised by the cre and thus will not interact with any subsequently added lox sites.

There have also been previous studies using random integration of lox sites to create deletions within a bacterial genome. Multiple random deletions were observed in *Corynebacterium glutamicum* using a similar method, with loxP sites contained within transposons and randomly integrated into the genome and activated via a cre suicide vector, ranging from 400 bp to 158 Kb in size (Tsuge et al., 2007). However, the system was not self selective, with the authors commenting that only 1.5% of final colonies contained deletions. The rest of the colonies contained inversions, and only 42 unique deletions were characterised. The authors also note that of the 42 deletion strains they recovered, only 2 had growth rates stronger than the WT strain, and many showed severe fitness defects. It is reasonable to assume that there may have been other deletion strains that could not compete against the large number of cells that had inversions, and therefore increased fitness due to no gene loss, and thus were lost from the pool.

By ensuring that our system leads to full selection against inversions between the lox sites, we can attempt to prevent this issue of competition against quasi-WT cells, and thus potentially retain as many different deletions as possible. By looking at them read numbers of each unique deletion, we can also get an estimate of how many cells were in the population, and thus an estimation of the growth rate compared to the other clones.

This study also highlighted the utility of using the cre recombinase as its own self-selective marker. The inversion between a left-mutant and right-mutant lox sites necessitated the creation of a loxP site (Ghosh and Van Duyne, 2002; Van Duyne, 2015), and thus we propose the action of the cre alone was enough to cause this self selection and retention on only the cells that contained a deletions resulting in an inactivated lox72. We have previously shown that a lethal phenotype is expressed in *M. pneumoniae* when the cre recombinase acts upon a lone active lox site (Shaw, 2019), and this worked perfectly as a counter-selective method here. The one caveat to this method being that 50% of potential deletions were removed due to the formation of the loxP instead of the lox72. However, looking at the overall distribution of deletions in Figure 4A, it is clear that the vast majority of putatively NE regions were deleted somewhere within our pool. Therefore, the high transposons density appears to be able to counter-balanced this loss in efficiency. A further improvement could be to use PacBio or Oxford Nanopore deep-sequencing technologies to ensure we reach the inverted repeats of both transposons and therefore we could map the deletions to a single base resolution.

Due to this high transposon density in the transformations, we achieved a very high coverage of deletions across the *M. pneumoniae* genome, resulting in the deletion of 21% of the genome across the pool, with deletions ranging from under 50 bp to 28 Kb. However, the distribution of these deletions is not uniform, with the six most represented deletions accounting for over 99% of the dataset. Whether this is a true reflection of the ratio of deletion mutants, or an unforeseen by-product of the sequencing and analysis pipeline is unclear. Despite this, we were able to validate deletions with low read numbers. Regions B and C consisted of 0.047% and 0.015% of the total reads respectively and were easily identified, and region D was isolated via PCR and sequenced, despite accounting for only a single read. This indicates that the full list of 285 deletions is likely reliable. It also indicates that there was probably a bias within the library preparation for the sequencing protocol that artificially inflated the most common deletions, thus skewing the final deletion ratio.

The high levels of variation within the larger regions also indicates the pool is robust and contains multiple valid deletions, as it shows that the variation of insertion sites for the original lox site transposons is as high as we expected. The concentration of deletions in hotspots is not due to a bottleneck caused by poor transformation efficiency in either of the lox insertion stages, as the variation in the hotspots shows multiple integrations, with a vast range in the size of the deletions across the general region. If the hotspot was caused by the fact that only a small number of transposons were present in one of stages, the vast majority of the deletions would share a common end or starting point, which is not what we observe. Instead, we are probably seeing those regions whose loss imparts the lowest reduction in cellular fitness. Looking at the region between bases 529000 and 630000 (Indicated by the blue bars in Figure 4A), we see a very high density of deletions. This region is almost entirely populated by NE genes (24 NE genes, 3 F genes), of which 17 have no clearly defined function (see Suppl. Table 7, *mpn490* to *mpn513*).

Due to the uncertainty inherent to the data generated from the sequencing protocol, i.e. the majority of reads not containing both inverted repeat regions, we decided that splitting the genome into 50 bp bins gave us specificity enough to map the deletions as accurately as possible. Due to this aggregation method however, there could be many more deletions that are similar to each other by fewer than 50 bp, and thus are missed from the analysis by being grouped with the other reads. This could mean however that we are greatly underestimating the number of unique deletion events that occured.

A major consideration with this protocol is its population based, and thus competitive nature. The large variation of deletions that are created are growing in direct competition with each other, and thus this protocol also allows for us to partially select for those cells that are the most robust. While we only allowed for one passage for the cells to grow in an attempt to minimise this as much as possible, the selection for faster growing mutants is inevitable, and it remains an inherent property of bacterial life that the fast-growers will proliferate at the expense of the slow growers. One possibility to have a larger coverage could be to plate the cells in agar and once grown scrape all and sequence. This way since cells will grow as single colonies competition will be minimized.

By utilising a randomised large scale deletion protocol, not only can single genes be tested for their updated esentiality, but whole regions as well. This has the power to greatly increase the scope of genome reduction projects, by quickly identifying the largest areas that are amenable to deletion at any given time, and identifying those regions that may not be amenable to deletion despite being counterintuitive. This is shown well by the deletions we see across the population. While the distribution of E and NE genes is fairly even in the *M. pneumoniae* genome, there are some clear islands of non-essentiality. We have shown that large deletions are possible in many of these islands, the regions around the 610 Kb and 140 Kb most notably. Looking at Figure 4A, the majority of the regions with multiple NE genes contain some level of deletion. However, there are also places where few if any deletions are observed, such as the clusters of NE genes at 244 Kb, 370 Kb and 490 Kb. While the loss of any of these genes individually is possible for the cell, their combined loss appears to be lethal. Therefore, this tool can be used not only to find which regions of the genome are most amenable to large scale deletion, but also to identify which putative nonessential regions play host to the most epistatic interactions.

Furthermore, the competitive nature of the protocol allows the researcher to not only elucidate the larger regions that can be deleted, but also those with the least fitness expense in any given environment. Similar to the RANDEL protocol outlined by Vernyik et al, we can place emphasis on cell fitness and robustness of growth during selection (Vernyik et al., 2020). As LoxTnSeq does not rely on endogenous DNA repair mechanisms for it’s deletion however, it has the potential to be applicable in a wider range of organisms.

If a cell is being minimised for a specific application, then genome reduced pools can be grown in the desired condition for as long as required, and the cells with the most viable reductions will outcompete those whose deletions are less viable, and give researchers insights into not only which genes provide a desirable phenotype in new conditions, but which operons and larger genome areas as well. We found limited examples of this in our own study. Our pool of deletions was grown for just a single passage under standard laboratory conditions, and thus genes required for the *M. pneumoniae* cells to exhibit pathogenesis were not required. As such, 15 separate adhesion proteins (out of a total of 22) were among those deleted, as were nine restriction putative enzyme proteins. On top of this, the main virulence factor in *M. pneumoniae*, the CARDS toxin (Parrott et al., 2016; Waites and Talkington, 2004), was also among those genes deleted, along with many of the adhesins that are linked to pathogenesis (Parrott et al., 2016). This indicates that the protocol has the ability to be utilised as an attenuation process as well.

In line with this, the methodology also has potential for conversion into a multi-step protocol. The removal of the constitutive cre transposon from the genome, or its replacement with a conditionally activated cre could allow for multiple rounds of the technique to be utilised within a single cell. This could convert the technique from a screening tool to identify amenable deletions to a self contained large scale genome minimisation technique, capable of deleting as many large and small genomic regions as the cell can endure. Coupled with it’s random nature, the technique could help researchers avoid unforeseen negative epistatic interactions and delete as much genetic material as is feasible within the cell, potentially paving the way for further elucidation of the minimal machinery needed for a cell to survive. While single rounds of the protocol may only remove relatively small amounts of DNA in this case (our largest deletion of 28 Kb accounts for 3% of the total *M. pneumoniae* genome), its unbiased and competitive nature make LoxTnSeq an attractive prospect.

In conclusion, we present the LoxTnSeq protocol as a multi-purpose tool for synthetic biology. It is capable of deleting large regions of NE genes from a host genome, identifying candidate regions for genome reductions based on the fitness the deletion imparts on the cell, allowing for more accurate essentiality maps based on the loss of multiple genes. It also allows for the identification of NE regions that contain strong epistatic interactions that may cause a loss of viability if removed together, despite their constituent parts being deemed NE. We hope that these attributes will make it a useful contribution to the growing synthetic biology tool box.

## Methods

### Strains and culture conditions

Wild-type *Mycoplasma pneumoniae* strain M129 (ATTC 29342, subtype 1, broth passage no. 35) was used. Cells were cultured in 75cm^2^ tissue culture flasks at 37°C in standard Hayflick media, as described by Hayflick (1965) and Yus et al., (2009) (Hayflick, 1965; Yus et al., 2009), supplemented with 100 μg/ml ampicillin, 2 μg/ml tetracycline, 20 μg/ml chloramphenicol, 3.3 μg/ml puromycin and 200 μg/ml gentamicin as appropriate. Hayflick agar plates were created by supplementing the Hayflick with 1% Bacto Agar (BD, Cat. No. 214010) before autoclaving.

NEB 5-alpha Competent *E. coli* cells (New England Biolabs, Catalogue number C2987H) were used for plasmid amplification and cloning. They were grown at 37°C in Lysogeny Broth (LB) at 200RPM or static on LB agar plates, supplemented with 100 μg/ml ampicillin.

### Plasmid DNA

All plasmids were generated using the Gibson isothermal assembly method (Gibson et al., 2009). DNA was isolated from NEB 5-alpha Competent *E. coli* cells, and individual clones were selected by using LB + ampicillin plates (100 μg/ml). Correct ligation was confirmed via Sanger sequencing (Eurofins Genomics). A list of all plasmids, and primers used in their generation and sequencing can be found in Suppl. Tables 9 & 10. The plasmids used in this study were generated as follows:

#### pMTnLox71Tc

The plasmid pMTnTetM438 was amplified separately via PCR using oligos 1 & 2, and 3 & 4. The samples were digested with DpnI and isolated via electrophoresis. Bands of approx. 4.2 Kb and 2 Kb respectively were isolated and annealed via Gibson assembly.

#### pMTnLox66Cm

The plasmid pMTnCat was amplified via PCR using oligos 5 & 6. The sample was digested with DpnI and isolated via electrophoresis. A band of approx. 5 Kb was isolated and self-annealed via Gibson assembly.

#### pMTnCreGm

The plasmid pMTnCat was amplified via PCR using oligos 7 & 8, and the plasmid pGmRCre was amplified via PCR using oligos 9 & 10. The samples were digested with DpnI and isolated via electrophoresis. Bands of approx. 4.2 Kb and 2.6 Kb respectively were isolated and annealed via Gibson assembly.

### Random genome reduction protocol

WT M129 cells were transformed according to the protocol outlined by Hedreyda et al (Hedreyda et al., 1993). Cells were grown to mid-log phase, identified by the Hayflick media changing from red to orange. The media was decanted and the flask was washed 3x with 10 ml chilled electroporation buffer (EB: 8mM HEPES, 272nM sucrose, pH 7.4). Cells were scraped into 500 μl chilled EB and homogenised via 10x passages through a 25-gauge syringe needle. Aliquots of 50 μl of the homogenised cells were mixed with a pre-chilled 30 μl EB solution containing the 1 pMole of pMTnLox66Cm plasmid DNA. Samples were then kept on ice for 15 mins. Electroporation was done using a Bio-Rad Gene Pulser set to 1250 V, 25 μF and 100Ω. After electroporation, cells were incubated on ice for 15 mins, then recovered into a total of 500 μl Hayflick media and incubated at 37°C for 4 hours. 125 μl of transformed cells were then inoculated into T75 cm^2^ culture flasks containing 20 ml Hayflick and supplemented with 20 μg/ml chloramphenicol.

The transformed cells were grown to mid-log phase, then isolated via pelleting. The described protocol was replicated in duplicate, one of the samples was sequenced to validate the transposon insertion using pMTnLox66Cm (R1B1) while the second was used in the subsequent transformations (R1B3). To ensure the recovery of planktonic cells, the media was transferred to a 50 ml Falcon tube. The flask was then scraped into 500 μl Hayflick, which was added to the Falcon tube with the media. The sample was centrifuged at 10,000 RPM at 4°C for 10 mins to pellet the cells. The supernatant was discarded and the cells resuspended in 500 μl chilled EB. The cells were homogenised via 10x passages through a 25-gauge syringe needle. Aliquots of 50 μl of the homogenised cells were mixed with a pre-chilled 30 μl EB solution containing the 1 pMole of pMTnLox71Tc plasmid DNA, and transformed using the previously described settings. Cells were recovered in the same manner, and 125μl were inoculated into a T75 cm^2^ flask containing 20 ml Hayflick media supplemented with 2 μg/ml tetracycline and 20 μg/ml chloramphenicol.

The transformed cells were again grown to mid-log phase and isolated via the centrifugation method described above. Aliquots of 50 μl of the homogenised cells were mixed with a prechilled 30 μl EB solution containing the 1 pMole of pMTnCreGm plasmid DNA, and transformed using the previously described settings. Cells were recovered in the same manner, and 125 μl were inoculated into a T75 cm^2^ flask containing 20 ml Hayflick media supplemented with 200 μg/ml gentamicin.

### Quantification of transposon density

Cultures R1B1 (transformed only with pMTnLox66Cm) and R1B3 (R1B1 transformed with pMTnLox71Tc) were grown to mid-log phase then cells were isolated via the centrifugation protocol outlined above. Genomic DNA was isolated via the MasterPure™ genomic DNA purification kit and sent for sequencing using a standard 125 bp paired-end read library preparation protocol for an Illumina MiSeq. The raw data was submitted to the ArrayExpress database (http://www.ebi.ac.uk/arrayexpress) and assigned the accession identifier E-MTAB-9590. Insertion sites for the transposon were identified using Oligo 19 (Suppl. Table 10), which is bound to the 3’ end of the chloramphenicol resistance genes, directly downstream of the inverted repeat (IR) sequence, and identified in the *M. pneumoniae* M129 genome using FASTQINS (Miravet-Verde et al., 2020). This bioinformatic tool allows to identify the point of insertion of the transposon used in the study and quantify, by read counts, how many times that specific insertion is detected by sequencing. This tool was run in both samples R1B1 and R1B3, corresponding to the population transformed with pMTnLox66Cm, and the secondly transformed with pMTnLox71Tc, respectively (Suppl. Table 1). These results were later analysed using ANUBIS essentiality framework (Miravet-Verde et al., 2020) to explore the coverage (i.e. percentage of nucleotide bases found inserted for a considered set of positions), considering the genome of *M. pneumoniae* (816,394 bp) and for each set of known essential and non-essential genes as described by Lluch Senar et al. (Lluch-Senar et al., 2015). For the analysis of the location of the insertions, we considered the gene density (i.e. number of insertions normalized by the gene length, Suppl. Table 4) and the genome of *M. pneumoniae* at 1kb resolution (bin size of 820bp, Suppl. Table 2). Pearson’s R correlations can be found in Suppl. Table 3.

### High throughput deletion sequencing

From the original isolated cells that survived transformation with the pMTnCreGm, 100μl was isolated, and had genetic material isolated via the MasterPure™ genomic DNA purification kit (Lucigen, Cat. No. MC85200). Genomic DNA was fragmented to 300 bp via Covaris sonication. 5’ phosphorylation was undertaken to allow for adapter binding, then 3’ overhangs were filled to create blunt ends. These were then ligated using T4 ligase to create circular fragments, and linear DNA was removed via digest with exonuclease I and lambda exonuclease. Circular DNA was then denatured and amplified using an oligo mix containing oligos 20, 21, 22 & 23, and a phi29 polymerase to amplify DNA containing a deletion scar. This amplified product was then fragmented again using Covaris to 300 bp and NEBNext Adaptor for Illumina were annealed to the linearised DNA. This DNA was then sequenced using paired end reads of 150 bp in an Illumina Hi-Seq 2500. The raw data was submitted to the ArrayExpress database (http://www.ebi.ac.uk/arrayexpress) and assigned the accession identifier E-MTAB-9582.

From the sequencing data, paired reads were extracted that contained the inverted repeat sequence from the transposons, a sequence of genomic DNA, the adapter sequence from the circularisation protocol and a second sequence of genomic DNA. We used basic bash tools to trim the adapter sequence and BlastN to map against *M. pneumoniae* M129 genome (Accession number: NC_000912). Then we used custom Python scripts to detect deletion points selecting for reads where the two halves of the read map to two different genomic loci. The scripts required to run these processing steps can be found in https://github.com/CRG-CNAG/Fastq2LoxDel. With this we extracted a list of genomic positions representing each side of a deletion and a read count value representing how many times they were found (Suppl. Table 1).

### Validation of inversion removal

To ensure that the protocol only selected for cells that had undergone a deletion, the pool of cells containing all three transformations was grown to mid-log phase and isolated via the centrifugation protocol described above. The cells were then serial diluted in Hayflick media to a 10^-5^ concentration, and 100 μl was lawn plated onto freshly prepared Hayflick agar plates. Plates were incubated at 37°C for 10 days. 100 individual colonies were picked and suspended in wells containing 200 μl of Hayflick in 96 well plates. The cultures were homogenised, and separate 50 μl aliquots were inoculated into wells containing 150 μl Hayflick supplemented with 25 ng/μl chloramphenicol, 150 μl Hayflick with 2.5 ng/μl tetracycline and 150 μl plain Hayflick respectively. 150 μl Hayflick was also added to the original well to restore it to 200 μl. The plates were incubated at 37°C for 7 days. After this time, colonies were assayed for their ability to grow in plain Hayflick media vs media containing the antibiotic resistances conferred by the transposons.

### Deletion validation

Selected deletions uncovered through the sequencing protocol were validated using PCRs of the genomic DNA from the pooled cells. 100μl of the cells that grew after transformation with the pMTnCreGm plasmid had their genomic DNA isolated via the MasterPure Genomic DNA Extraction Kit (Invitrogen). Oligos described in Suppl. Table 9 were used to amplify specific regions in both the deletion pool and WT DNA, and differences in band sizes were visualized via gel electrophoresis. Specific bands showing a deletion were cut and DNA was purified via the QIAGEN Gel Purification Kit and sequenced via Sanger sequencing to confirm the presence of the lox72 site and genomic loci.

---

Figure S1: Frequency of deletions by size in our database, with all genes containing no essential genes in light green, and all deletions containing an essential gene in blue.

