## Supplementary figures and images for "LoxTnSeq: Random Transposon insertions combined with cre/lox recombination and counterselection to generate large random genome reductions"

### Suppl.Figure 1

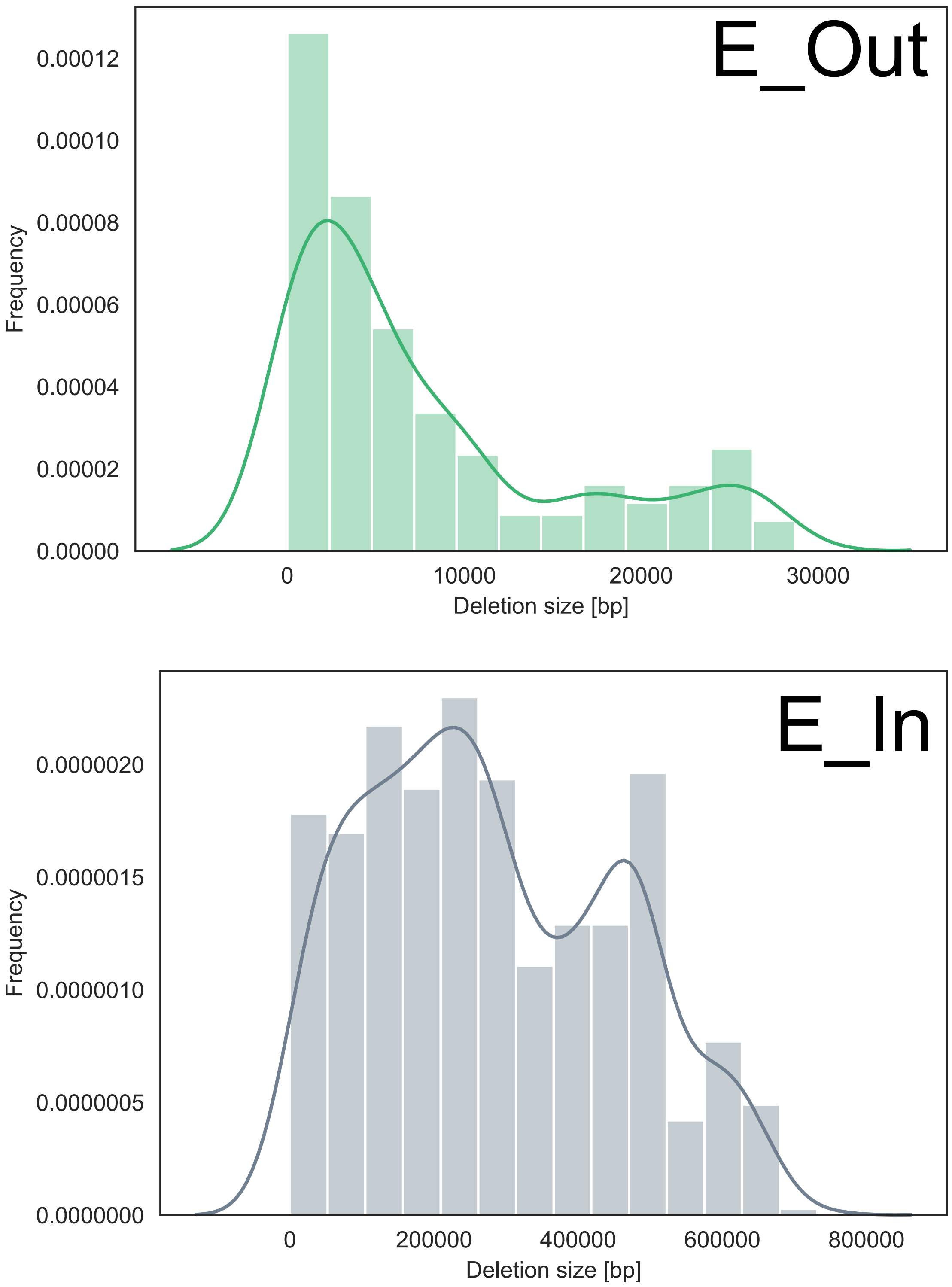
